# Evolutionary responses to codon usage of horizontally transferred genes in *Pseudomonas aeruginosa*

**DOI:** 10.1101/2020.07.11.198432

**Authors:** Martijn Callens, Celine Scornavacca, Stéphanie Bedhomme

## Abstract

Prokaryote genome evolution is characterized by the frequent gain of genes through horizontal gene transfer (HGT). For a gene, being horizontally transferred can represent a strong change in its genomic and physiological context. If the codon usage of a transferred gene deviates from that of the receiving organism, the fitness benefits it provides can be reduced due to a mismatch with the expression machinery. Consequently, transferred genes with a deviating codon usage can be selected against or elicit evolutionary responses that enhance their integration. In this study, a comparative genomics approach was used to investigate evolutionary responses after the horizontal transfer of genes with diverse degrees of codon usage mismatch in *Pseudomonas aeruginosa*. Selection on codon usage of genes acquired through HGT was observed, with the overall codon usage converging towards that of the core genome over evolutionary time. This pattern seemed to be mainly driven by selective retention of transferred genes with an initial codon usage similar to that of the core genes. Gene amelioration, through the accumulation of synonymous mutations after HGT, did not seem to systematically affect transferred genes. Additionally, variation in the copy number of tRNA genes was often associated with the acquisition of genes for which the observed variation could enhance their expression. This provides evidence that compensatory evolution might be an important mechanism for the integration of horizontally transferred genes.

## Introduction

Horizontal gene transfer (HGT) strongly contributes to bacterial evolution and is a key mechanism for a quick adaptation to environmental challenges. The frequent uptake of foreign genetic material leads to numerous genes being found in certain genomes of a bacterial species, but not in all. This set of dispersed genes, the accessory genome, is highly dynamic and is shaped by frequent gene gains and losses (Vos et al. 2015). A new gene is incorporated into the accessory genome if – after initial acquisition – it replicates itself within the receiving organism and is vertically inherited. However, long-term retention of a gene will depend on its effect on the fitness of the receiving organism, which is often dependent on the environmental and genomic context. For a gene to be retained over evolutionary time, the cost of replicating the transferred genetic material and expressing any genes contained in it has to be counterbalanced by a net fitness benefit. Otherwise, there is a high probability that it will be subsequently lost due to natural selection (Koskiniemi et al. 2012).

One factor that can influence the cost of expressing a horizontally acquired gene is the degree of match of its codon usage to the translation machinery of the organism expressing it (Baltrus 2013). Within a species, synonymous codons are often not used at equal frequencies but particular codons are preferred over synonymous alternatives to encode a same amino acid (Grantham et al. 1980). These codon usage preferences are a well-documented phenomenon in bacteria and can be strongly differentiated between species (Novoa et al. 2019). Although neutral processes can shape codon usage bias due to a species-specific mutational bias, in unicellular organisms with large effective population sizes there is often a selective component involved (Hershberg and Petrov 2008). In bacteria there is usually a good correlation between tRNA gene copy number and the frequency of corresponding codons, and preferred codons are generally more efficiently and accurately translated than less frequently used ones because of a higher degree of match to the available tRNA pool (Rocha 2004; Higgs and Ran 2008). Consequentially, genes using rare codons often produce a low amount of functional protein due to slower translation, translation errors, protein truncation or protein misfolding (Plotkin and Kudla 2011). Furthermore, mRNAs containing non-optimal codons are found to be less stable than their synonymous equivalents using optimal codons (Presnyak et al. 2015). Sequestration of ribosomes on the mRNA is also higher when elongation rates are slow, reducing ribosome availability and global protein synthesis (Kudla et al. 2009; Frumkin et al. 2018).

Horizontally acquired genes with a codon usage that is divergent from the receiving organism will thus potentially have lower relative fitness effects than when codon usage is similar. Previous analyses have shown that the fixation probability is higher for genes with a codon usage that matches that of the receiving organism (Medrano-Soto et al. 2004), and is related to the capacity to translate these genes (Tuller et al. 2011). Moreover, if a gene with a divergent codon usage is retained after HGT, it is expected to evolve towards a more typical codon usage in a process called *amelioration* (Lawrence and Ochman 1997). This process relies on synonymous mutations accumulating in the transferred gene, and is assumed to be very slow when it is only driven by the local mutational bias. However, if selection on codon usage is involved this process can occur faster and will depend on the fitness effects of individual synonymous mutations.

Another evolutionary mechanism to accommodate for a transferred gene with divergent codon usage is *compensatory evolution* (i.e. mutations elsewhere in the genome that reduce the cost associated to the transferred gene; Bedhomme et al. 2019). This mechanism has been shown to play an important role in preventing plasmid loss and the fixation of accessory traits in the chromosome (Harrison et al. 2016). A potential strategy for compensatory evolution to alleviate the cost of divergent codon usage in transferred genes is increasing the availability of tRNAs that recognize the rare codons they contain (Tuller 2011). One way to do this is by increasing the gene copy number of these tRNAs, which is known to be correlated with cellular tRNA concentrations (Dong et al. 1996). In bacteria, there are indications that this strategy is used to compensate for HGT-related changes in codon usage (McDonald et al. 2015).

*Pseudomonas aeruginosa* is an ecologically versatile opportunistic pathogen that frequently acquires genes through HGT. This is reflected by the large number of genes found in its accessory genome, which are often acquired as genomic islands (i.e. functionally related clusters of genes inserted at specific genomic locations). Important phenotypic traits displayed by some strains, such as increased pathogenicity, are known to be related to the acquisition of genomic islands through HGT (Harrison et al. 2010). In this study, we investigate the extent of gene acquisition through HGT in *P. aeruginosa* and the subsequent evolutionary responses related to codon usage in transferred genes. To this aim, we applied a comparative genomics approach using full genome sequences of different isolates of *P. aeruginosa* and addressed three questions: 1) Is there an initial selective retention of genes after HGT based on their codon usage? 2) Does codon usage in horizontally transferred genes converge towards the codon usage of the receiving organism over evolutionary time, and is this driven by mutational processes? 3) Is tRNA gene copy number variation correlated with HGT-related changes in codon usage?

## Results

### Pan-genome analysis, strain phylogeny and core genome codon usage

A schematic overview of the bioinformatic pipeline used in this study is provided in Figure S1. For pan-genome determination of our collection of 94 *P. aeruginosa* genomes, homologous genes were identified by sequence clustering on the complete set of protein coding genes. Homologs present in more than 88 strains were assigned as core genes, while the remaining genes were considered to be accessory genes. This analysis gave a total of 16 007 distinct clusters of homologous genes. From them, 5288 were classified as core genes, based on the numbers of strains they occurred in (4330 genes occurred in all 94 strains and 958 genes occurred in between 89 to 93 strains). A total of 10 719 homologous gene clusters were present in less than 89 strains and were classified as accessory genes.

We constructed a phylogeny of the *P. aeruginosa* strains used in this study based on the nucleotide sequence alignment of their core genes. This phylogeny indicated a structure consisting of two large clades, with 61 strains belonging to the clade containing reference strain PAO1 and 32 strains belonging to the clade containing reference strain PA14 (Figure 1). Strain PA154197 branched off close to the root and did not seem to be closely related to either one of these two clades. Although bootstrapping indicated a good support for this general structure, some smaller clades were found to be less -well supported (Figure S2).

**Figure 1:**
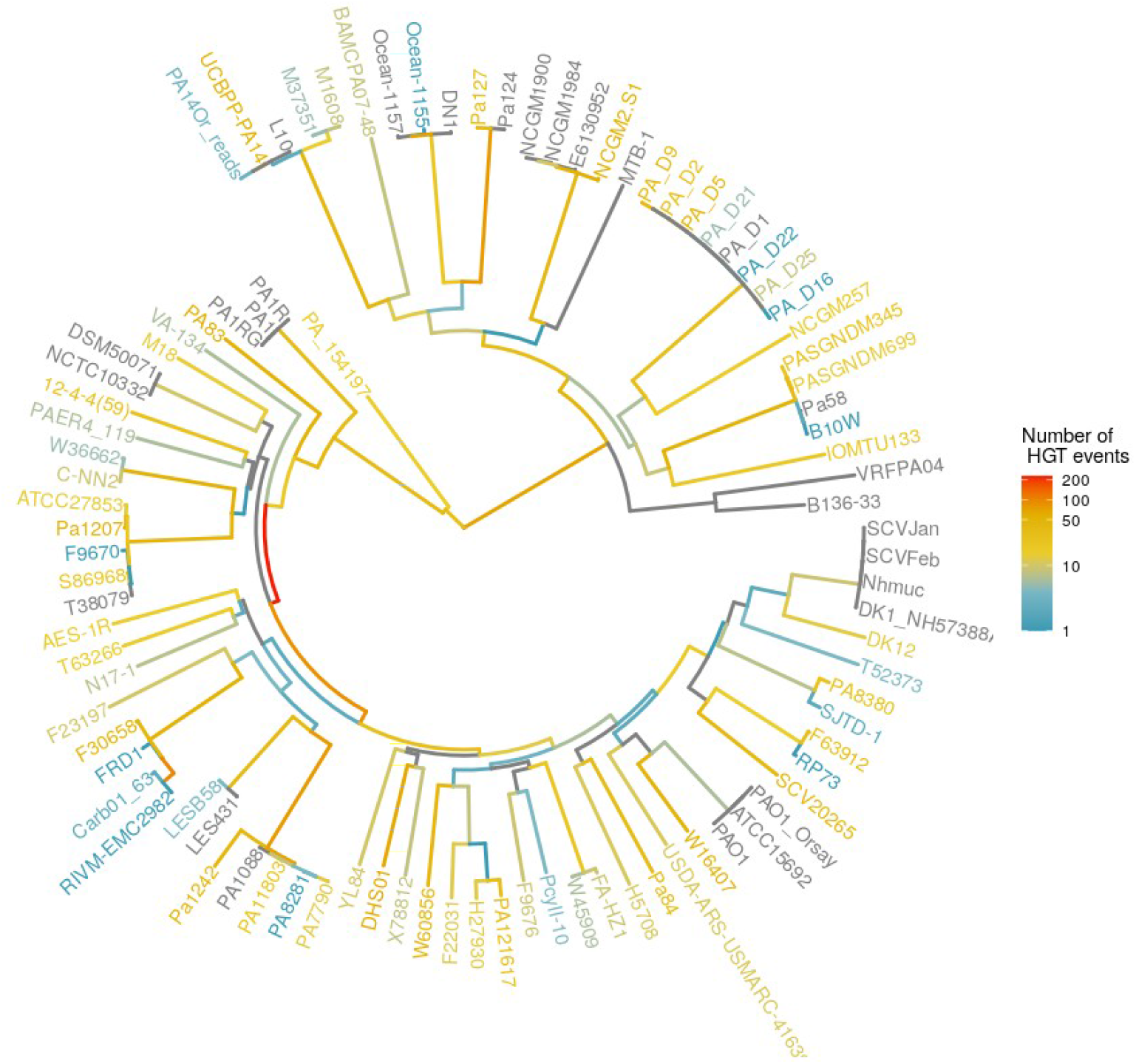
Number of HGT events along the evolutionary history of *P. aeruginosa*. The phylogeny of *P. aeruginosa* was determined using the alignment of the core genes. Branch color indicates the number of HGT events detected along each branch. The color of the tip labels give an indication of the number of HGT genes that were unique to a strain.

Analysis of the GC-content of the core genome gave an average of 67.4 %GC for coding regions and 62.0 %GC for intergenic regions. The GC-content of full genomes was slightly lower than the core genome (66.4 %GC for coding regions and 60.7 %GC for intergenic regions). The construction of a codon usage table for the core genes showed that the codon usage in core genes reflected this relatively high %GC, where G or C nucleotides were always favored over A or T on degenerate first and third codon positions (Table S1). We calculated ENC values (Wright 1990) for the core genes. This metric indicates the strength of codon usage bias in a gene, and ranges between 20 (only one codon is used for encoding each amino acid) and 61 (all synonymous codons are used more or less equally). Core genes generally had a strong codon usage bias, as shown by their overall low ENC values (mean = 30.8 ± 3.8; Figure 2C).

**Figure 2:**
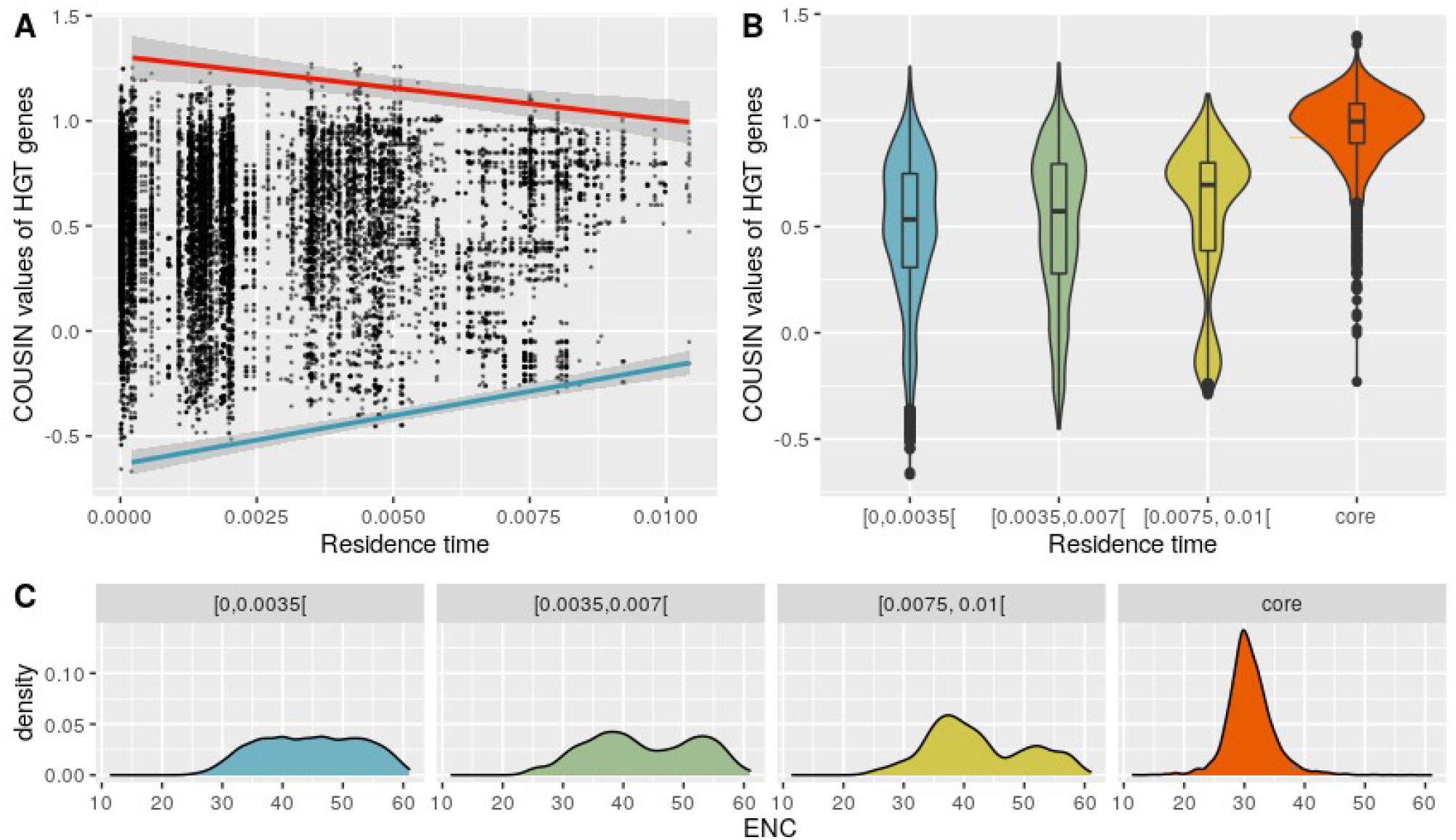
Codon usage bias of HGT genes in relation to their residence time. A: Scatterplot of the COUSIN value of HGT genes in function of their residence time. The blue and red regression lines were determined by taking respectively the minimal and maximal COUSIN value for each residence time interval of 0.001. B: Violin plot showing the distribution of COUSIN values of HGT genes for each residence time interval of 0.0035 and the core genes. C: Density plots showing the distribution of the effective number of codons for HGT genes for each residence time interval of 0.035 and for the core genes.

### Inference of horizontal gene transfers

We compared accessory gene clusters to all other non-homologous gene clusters to filter out clusters that were classified as accessory genes due to erroneous classification to a separate cluster. With this analysis we identified a large fraction of accessory genes that showed evidence of pseudogenization (33.4% of all accessory genes) based on their low sequence overlap (<70%) but high shared sequence identity (>90%) with non-homologous gene clusters. One-third of these putative pseudogenes were partial sequences of larger genes that occurred in more than 89 strains, and could thus be considered truncated versions of duplicated core genes (11.5% of all accessory genes). The remaining putative pseudogenes were mainly length variants of genes occurring in only a limited number of strains. However, a few cases involved genes with high similarity to non-homologous genes because of shared conserved domains (e.g. conserved domains in the phosphate-selective porin O and P family). Although these last cases could represented genuine non-homologous clusters and HGT’s, they were not retained in the further analysis because differences were often not clear-cut and we opted for a stringent filtering to avoid false positives.

After applying the previous filtering step, we performed an ancestral state reconstruction for all the remaining accessory genes. We first constructed phyletic patterns for each accessory gene on the strain phylogeny, with gene copy number as character states. The presence of an accessory gene was then inferred for each node on this phylogeny using ancestral state reconstruction with stochastic character mapping. This analysis inferred that 6.9% of the accessory genes were present in the common ancestor of all strains included in this dataset, indicating that these genes were classified as accessory genes because of multiple deletions throughout the evolutionary history of *P. aeruginosa*. Further evidence for multiple deletions of genes present in the common ancestor was provided by the large number of genes that were found at the same genomic location, but with a highly dispersed occurrence on the strain phylogeny (545 genes were present in only a small number of strains on both sides of the root at the same genomic location).

As in our analysis we were only concerned with accessory genes that originated through HGT, we excluded genes that were classified as accessory genes by potential misclassification or gene deletion. Based on these criteria, 6393 (59.6%) of the accessory genes were presumed to have been acquired by HGT.

### Horizontal gene transfers in *P. aeruginosa*

Horizontally transferred genes acquired along the same branch that were physically adjacent to each other were grouped and considered to represent a single HGT event. A total of 2990 individual HGT events were identified. The number of genes acquired per HGT event ranged from 1 to 37. Almost half of these events involved the acquisition of only a single gene, and 90% involved the acquisition of five genes or less. The number of HGT events along a single branch varied widely, even when correcting for branch length, and ranged from none to 230 (Figure 1).

The highest number of HGT events was recorded along the branch leading to the ancestral node of a clade containing 56 strains (230 events, 364 genes). A high number of HGT events was also recorded along the branch departing from this node, leading to a sub-clade of 44 strains (87 events, 146 genes). An exceptional high number of HGT events was further found along the branch leading to the common ancestor of strains Carb01_63 and RIVM-EMC2982, two multidrugresistant clinical strains isolated from different locations in the Netherlands (99 events, 188 genes). A total of 1165 HGT events, involving 4437 genes, were found to be strain-specific. Strain DHS01, a multidrug-resistant strain isolated in France, had the highest number of strain-specific HGT events (64 events involving 131 genes), although this number could be biased by the absence of closely related strains in our dataset.

Each gene acquired through HGT was assigned a *residence time*. This metric represents the evolutionary distance between the leaf on the phylogeny where a gene was found and the midpoint of the branch where this gene was acquired. A large number of genes acquired through HGT were characterized by a relatively short residence time, indicative of relatively recent transfers. The amount of genes strongly decreased with increasing residence time, with only a small number of HGT genes found in the pan-genome of *P. aeruginosa* that were acquired early in its evolutionary history (Figure S3).

### Codon usage of genes acquired by HGT

Genes acquired by HGT had a much lower average %GC than core genes (58.8 %GC and 67.4 %GC, respectively), and a larger variation, with values ranging from 30.6 %GC to 73.3 %GC. To quantify the codon usage bias of horizontally transferred genes, we used the Codon Usage Similarity Index (COUSIN; Bourret et al. 2019). This metric compares the codon usage of a gene against a reference codon usage table (the core genome codon usage table was used in our analysis). The values indicate both the direction of codon usage bias (positive values represent a bias in the same direction as the reference, negative values are in the opposite direction) and the strength (indicated by the absolute values, with 0 indicating the absence of a bias and values above 1 indicating a stronger bias than the reference). The majority of HGT genes had a codon usage bias in the same direction as the core genes (88.0% was characterized by a positive COUSIN value; Figure 2). The strength of codon usage bias of these genes was however more variable and generally lower than that observed for the core genes (mean ENC = 44.3 ± 8.7 and 30.8 ± 3.8, respectively; Figure 2C). HGT genes with a codon usage bias in the opposite direction of the core genome (COUSIN < 0) were generally also AT-biased (average 44.8 %GC) and showed an overall lower codon usage bias strength (mean ENC = 54.4 ± 4.3).

The overall relationship between residence time and codon usage was determined using a non-parametric correlation. A linear regression model was used to examine the relation between codon usage and the minimal, maximal and median values for residence time intervals of 0.001. There was an overall positive correlation between a gene’s COUSIN value and its residence time (Kendall’s rank correlation: τ = 0.027, p < 0.05). This positive correlation is mainly driven by an overall increase of the minimal COUSIN values in function of residence time (linear model for minimal values: p < 0.05; R^2^ = 0.93). The maximal COUSIN values showed a negative correlation with residence time (linear model for maximal values: p < 0.05; R^2^ = 0.63). We did not find a significant correlation between the median COUSIN values and residence time (linear model for median values: p = 0.1), indicating a converging trend of COUSIN values in function of residence time. The same trends were found for changes in CAI values in function of residence time (Figure S4) and for changes in COUSIN values in function of residence time when transfers near the root were not included (Kendall’s rank correlation: τ = 0.017, p < 0.05; linear model for minimal values: p < 0.05, R^2^ = 0.59; linear model for maximal values: p < 0.05, R^2^ = 0.58; linear model for median values: p = 0.3).

We further assessed the potential importance of gene amelioration for changes in codon usage in function of residence time (i.e. are genes evolving towards the codon usage bias of the receiving genome through mutational processes?). To evaluate this we inferred the ancestral sequences for HGT genes that were represented by at least 20 or more extant sequences in our dataset. Codon usage bias in extant and ancestral sequences were then compared to evaluate changes due to mutational processes. This analysis showed variable responses for different genes (Figure 3). Out of the 22 genes included in the analysis, 6 genes showed directional evolution towards the codon usage of the core genome, 10 genes showed directional evolution opposite to the codon usage of the core genome, and 6 genes did not show any sign of directional evolution. Table 1 shows that directional changes in codon usage bias were generally caused by both synonymous and non-synonymous variation. However, most directional changes remained significant when only taking synonymous variation into account.

**Figure 3:**
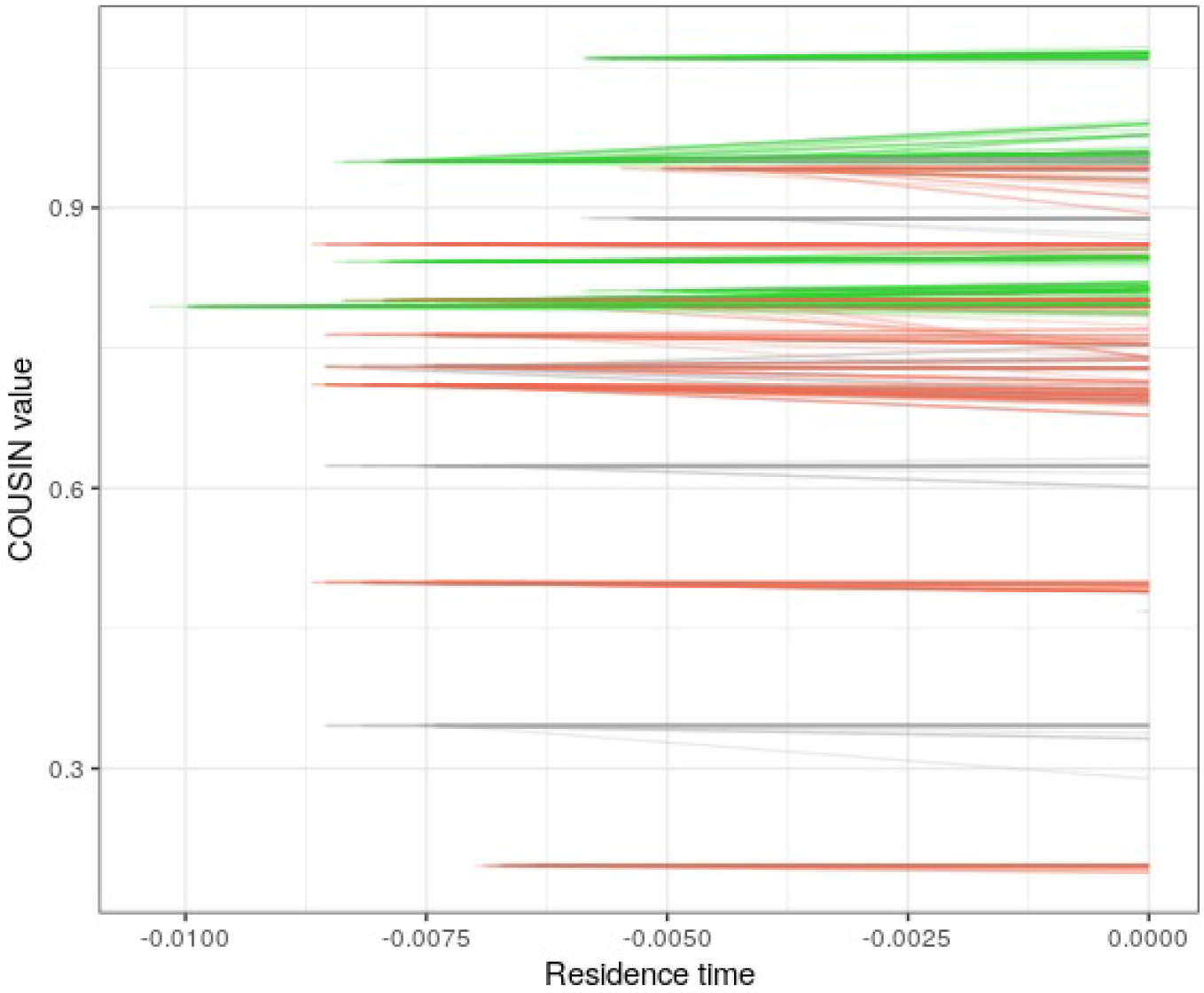
Changes in codon usage bias of HGT genes compared to their ancestral state. Each line represents the change in COUSIN value for a single CDS compared to its inferred ancestral sequence. Genes for which the mean change was positive are green, genes for which the mean change was negative are red and genes for which the change was non-significant are grey. Extant CDS’s were given residence time 0 and the ancestral sequence was given the value of the residence time of its corresponding CDS.

**Table 1:**
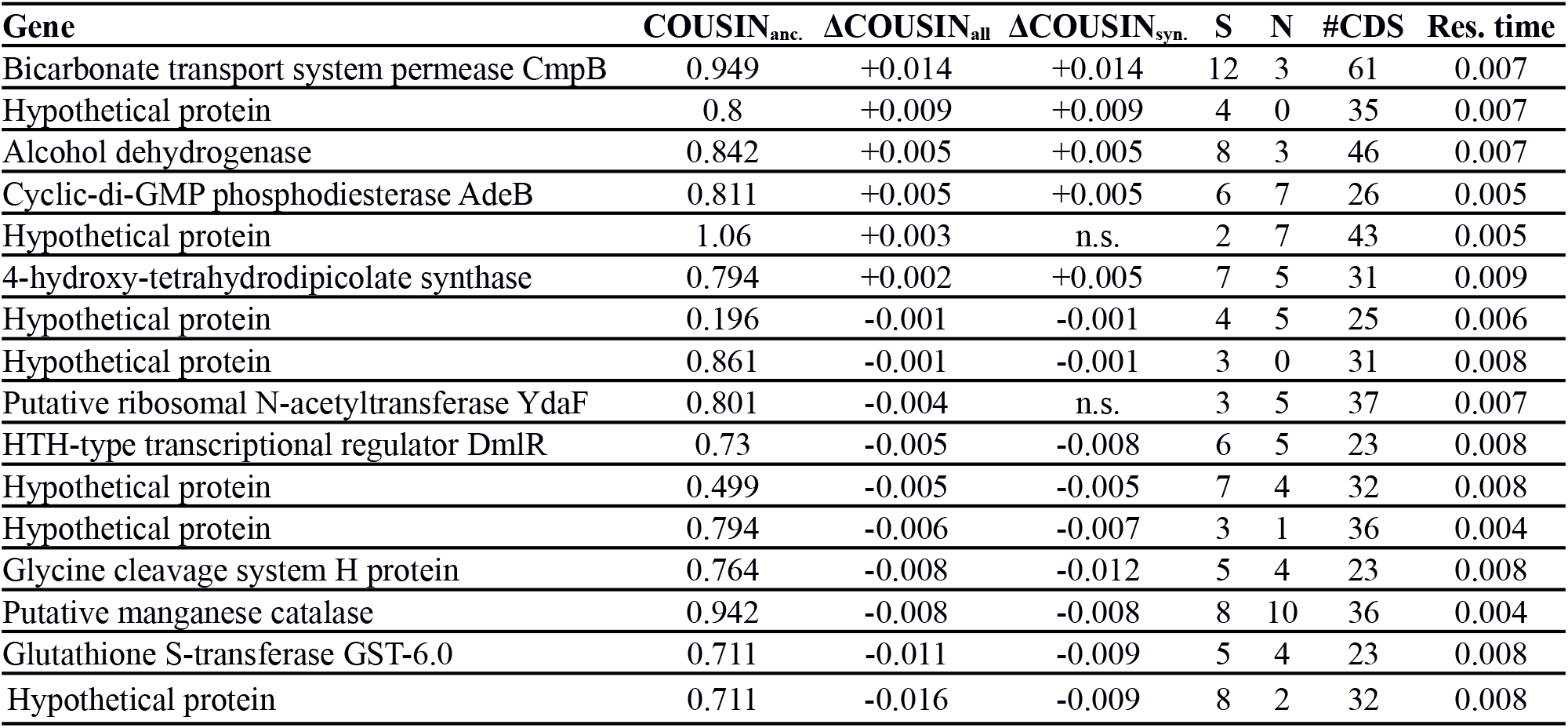
Genes with significant directional changes in their codon usage bias compared to their inferred ancestral sequence. COUSIN_anc._ = COUSIN value of the ancestral sequence; ΔCOUSIN_all_ = mean change in COUSIN values between ancestral and extant sequences. ΔCOUSIN_syn._ = mean change in COUSIN values between ancestral and extant sequences when only synonymous variation is taken into account. S = number of positions with synonymous variation. N = number of positions with non-synonymous variation. #CDS = number of CDS’s in the analysis for each gene. Res. time = mean residence time for each gene.

### tRNA gene copy number variation in relation to HGT

To reconstruct evolutionary changes in tRNA gene copy numbers, we first determined the gene copy number of each tRNA gene for each genome. We then inferred the gene copy number for internal nodes of the strain phylogeny using ancestral state reconstruction. A total of 98 events were detected involving a change in the tRNA gene content, with 83 of them being associated with the acquisition of genes through HGT. For most events this involved a change in the copy number of only a single tRNA gene (65 events), while some events showed variation in up to eight tRNA genes. Gene copy numbers of specific tRNA genes ranged from zero to eight copies, and 25 out of the 40 different tRNA genes found in *P. aeruginosa* had a variable gene copy number (Figure S5). The tRNA^Lys^_TTT_ gene, which had either two, three or four copies, showed the highest rate of changes in gene copy number, with 26 individual gene duplications and 44 individual gene deletions. The number of observed changes in gene copy number for other variable tRNA genes ranged between 1 and 13. There were overall more cases where the gene copy number of a specific tRNA increased than decreased (98 and 75 cases, respectively). For tRNA^Ala^_CGC_, a gene that is generally absent in *P. aeruginosa*, four independent gains were recorded, indicative of multiple acquisitions through HGT.

If variation in tRNA gene content provided compensatory evolution in response to the acquisition of genes with a divergent codon usage, we expected to observe two patterns. First, on a codon level, we expected that an increment in the copy number of tRNA genes would be associated with HGT events increasing the frequency of codons that these tRNAs can translate. Second, on a gene level, we expected that changes in tRNA gene content should have a proportionally larger positive effect on the translation of recently acquired HGT genes than on other genes.

We performed a permutation test to evaluate if increments in tRNA gene copy number were associated with strong increases in the frequency of codons these tRNA’s can translate. This analysis showed that there was a significant association with strong increases in the frequency of codons they can translate through Watson-Crick base pairing and wobble base pairing (p < 0.05 for both permutation tests), but not for less favored wobble base pairing interactions (Figure 4). Although most increases in the copy number of tRNA genes were found to be associated with increases in the corresponding Watson-Crick and potential wobble codons, for some tRNA genes we observed opposite response patterns. Increases in the copy number of tRNA^Ala^_CGC_ and tRNA^Arg^_CCG_ genes were never associated with increases in their corresponding codons. Also, increases in tRNA^Phe^_GAA_ gene copy number were always associated with an increased frequency of its wobble codon (TTT), while its corresponding Watson-Crick codon (TTC) always decreased in frequency.

**Figure 4:**
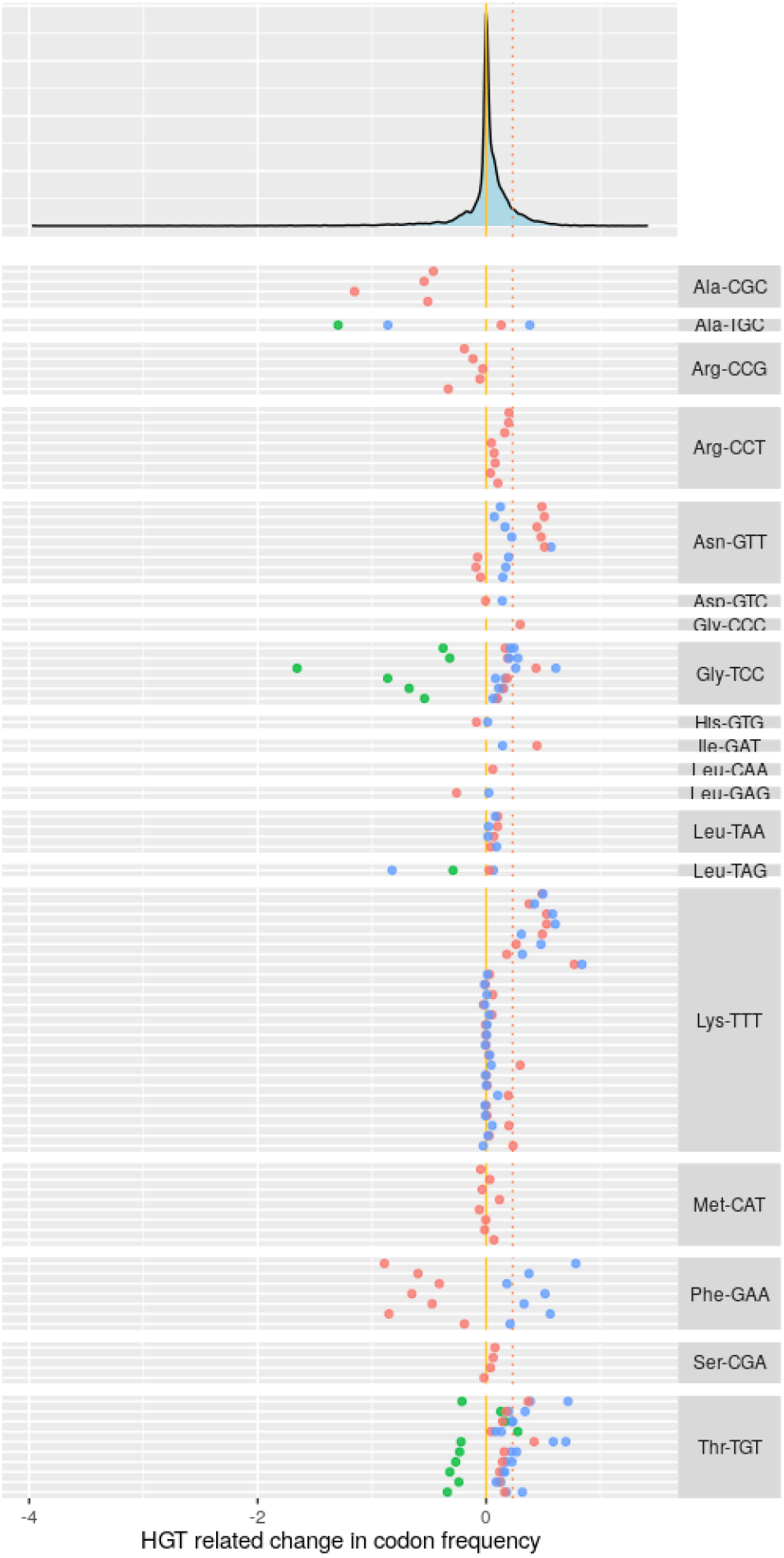
Changes in codon frequencies associated with increased tRNA gene copy numbers. Upper panel: Distribution of changes in codon frequencies along branches due to HGT (expressed as changes in the expected number of a codon per 1000 bases). The yellow line indicates no change in codon frequency, the red line indicates the 90^th^ percentile. Lower panel: Each horizontal line represents a single event of increase in tRNA gene copy number and are grouped per tRNA gene; Each point indicates a codon that was affected by this increase (red dots represent Watson-Crick base pairing, blue dots wobble base pairing and green dots less favored wobble base pairing). The position along the horizontal axis indicates the change in codon frequency due to HGT along the branch where the increase in tRNA gene copy number was observed.

To analyze compensatory evolution on a gene level, we used ΔtAI_abs_ values as a relative indication of how much a change in tRNA gene copy number is expected to enhance the translation of a gene. We calculated the ΔtAI_abs_ for genes acquired through HGT associated with a change in tRNA gene content that was expected to enhance overall translation. These ΔtAI_abs_ values were then compared to ΔtAI_abs_ values of genes that were present before the change in tRNA gene content occurred. We hypothesized that an expected increase in translation due to variation in tRNA gene content would be proportionally larger for recently acquired HGT genes than for other genes. In our dataset, there were 45 changes in tRNA gene content with an expected overall positive effect on translation, as determined by a mean positive ΔtAI_abs_ of all genes affected by this change. Of these, 29 changes in tRNA gene content caused a proportionally larger ΔtAI_abs_ for HGT genes than for genes that were already present at the parent node (Mann-Whitney U-test for each case: p < 0.05; Figure 5). In only 5 cases was the ΔtAI_abs_ higher for genes present at the parent node than for HGT genes (Mann-Whitney U-test for each case: p < 0.05), while in the remaining 11 cases there was no difference between the ΔtAI_abs_ of HGT genes and genes present at the parent node (Figure 5). On average, the increase in ΔtAI_abs_ was 1.5 times higher for HGT genes than for genes present at the parent node.

**Figure 5:**
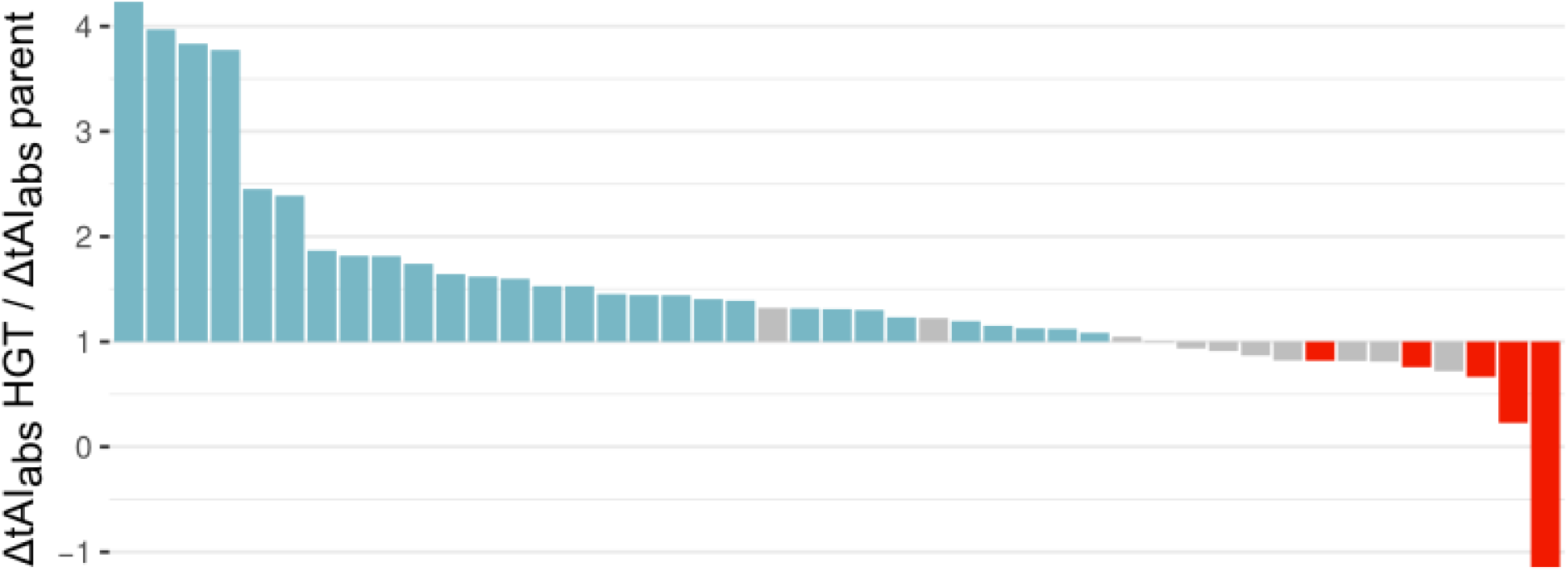
Change in ΔtAI_abs_ of horizontally transferred genes relative to the change in ΔtAI_abs_ of genes present at the parent node. Each bar represents an observed variation in tRNA gene content with an overall positive effect on translation. Values are obtained by dividing the expected change in translation efficiency (ΔtAI_abs_) for genes recently acquired through HGT and genes that were already present in each genome. Relative changes in ΔtAI_abs_ that were significantly larger for horizontally transferred genes are blue, those that were significantly larger for genes that were already present are red, non-significant differences are grey.

## Discussion

Evolutionary responses related to codon usage of horizontally transferred genes were investigated in *P. aeruginosa* using a comparative genomics approach. We showed that there was selection on codon usage of genes acquired through HGT, with their codon usage converging toward that of the core genome over evolutionary time. Gene amelioration, through the accumulation of synonymous mutations after HGT, did not seem to be driving the changes in codon usage towards that of the receiving organism. This favors the hypothesis that long-term retention of horizontally transferred genes was strongly influenced by their initial codon usage. We further found evidence that compensatory evolution might be important for the integration of horizontally transferred genes, as variation in tRNA gene content was often associated with the acquisition of genes for which the observed variation in tRNA gene content could enhance their expression.

Previous work already indicated that for multiple bacterial species the likelihood of successful HGT and subsequent gene retention increases when the codon usage of transferred genes is similar to that of the receiving organism (Medrano-Soto et al. 2004; Bolotin and Hershberg 2017). Our results show that this also plays a role in *P. aeruginosa*, especially for long-term gene retention. Initially there seems to be a relatively weak selection against the uptake of genes with an atypical codon usage. *P. aeruginosa* was found to incorporate genes exhibiting a wide range of codon usage, including unbiased genes and genes with a bias opposite to that of the core genes. However, the majority of recently transferred genes still had a codon usage bias in the same direction as the core genes, but generally less pronounced. Over evolutionary time, we observed selection against genes with a codon usage bias opposite to that of the core genes or with a very strong bias in the same direction as the core genes.

Two different mechanisms could drive the observed change in overall codon usage of genes acquired through HGT over evolutionary time. First, loss rates of genes with an atypical codon usage could be higher than for genes with a codon usage similar to that of the receiving organism. Especially if the maintenance and expression of genes with an atypical codon usage is costly, they are expected to be purged by natural selection (Koskiniemi et al. 2012). Even if the transferred genes do not confer a fitness cost, but are neither able to provide a net fitness benefit to the receiving organism due to their atypical codon usage, they will be prone to loss by genetic drift (Kuo et al. 2009). Second, gene amelioration due to the local mutational bias and/or selection on synonymous mutations could drive the codon usage of a transferred gene towards that of the receiving organism. We evaluated the potential role of gene amelioration and found no clear indications that synonymous mutations in horizontally transferred genes are causing the observed changes in codon usage. Although amelioration is assumed to equally affect all synonymous positions of non-preferred codons (Lawrence and Ochman 1997), it must be noted that with our approach parallel evolution (i.e. the occurrence of same synonymous mutations in different lineages) could cause an underestimation of the importance of this mechanism.

Amelioration is expected to occur when transferred genes are exposed to a new mutational and/or selective regime present in the receiving organism (Lawrence and Ochman 1997). The codon usage of the core genes is on the other hand assumed to be in equilibrium with these forces. There has been some speculation on the processes shaping the overall high GC-content and strong codon usage bias in the *P. aeruginosa* core genes. Dettman et al. (2016) experimentally showed that *P. aeruginosa* has a mutational bias towards AT. The same study noted that mismatch-repair (MMR) deficient strains had an inverse bias towards GC, and they hypothesized that a high prevalence of MMR deficient strains in combination with recombination would lead to a high equilibrium GC content. Furthermore, other mechanisms such as GC-biased gene conversion (Lassalle et al. 2015) or selection for high GC content (Hildebrand et al. 2010) could also play a role. The processes that are driving the high GC content and codon usage of the core genes did not seem to systematically induce synonymous variation in the same direction in horizontally transferred genes.

Although there was a general pattern of selection on codon usage towards that of the core genes, there were still several horizontally transferred genes with a long residence time that maintained an atypical codon usage bias. A notable example was the LESGI-7 genomic island (Figure 2: residence time ~ 0.008 and COUSIN < 0; Jani et al. 2016), which was inferred to be an ancient transfer at the base of the large clade containing strain PA14, and is shared by a large number of ecologically diverse strains. The 10 genes on this genomic island are involved in determining the lipopolysaccharide O-antigen serotype and exhibit a codon usage bias opposite to that of the core genes. A possible reason for the conservation of an atypical codon usage could be that gene expression is under the control of xenogeneic silencers that target DNA segments with a higher %AT than the chromosomal average (Singh et al. 2016). There are indications that xenogeneic silencing can act as a buffer against negative fitness consequences of HGT (Ali et al. 2014) and can facilitate the integration of horizontally acquired genes into existing regulatory networks (Will et al. 2015). In the *P. aeruginosa* strain PAO1, the xenogeneic silencer *MvaT* and its paralog *MvaU* are known to be critical for regulating the expression of virulence-associated and prophage genes (Castang and Dove 2012). It is highly probable that genes on LESGI-7, because of their high AT content, are targeted by the same xenogeneic silencing proteins. If regulation of genes on LESGI-7 is dependent on the interaction with these proteins, it would impose selection for maintaining a high AT-content to preserve regulation of expression and cause the conservation of their atypical codon usage.

We further found strong indications that the observed variation in tRNA gene content in *P. aeruginosa* was – at least partially – accommodating for the expression of horizontally transferred genes. Similar indications were found in *E. coli* using a comparative genomics approach (McDonald et al. 2015). In this species it has also been shown, through the experimental manipulation of tRNA gene copy number, that this mechanism can provide an efficient response to enhance the expression of genes with a codon usage that is not adapted to the tRNA pool (Du et al. 2017). Our results indicate that this mechanism might be more widespread in other bacterial species as a means for providing compensatory evolution related to HGT. Although we often found a strong association between variation in tRNA gene content and HGT events, there were also a large number of HGT events that caused the influx of atypical codons but were not associated with variation in tRNA gene content. Varying tRNA gene copy numbers is only one of several possibilities to compensate for the inefficient expression of genes with an atypical codon usage. Mechanisms such as the regulation of the expression of specific tRNA genes (Torrent et al. 2018), the action of tRNA modifying enzymes that can extend wobble base pairing (Endres et al. 2015) or mutations in the promoter of the horizontally transferred gene that increase its expression (Amorós-Moya et al. 2010) are other possibilities to improve the expression of horizontally transferred genes in their new host genome, but our analysis did not allow to test for these mechanisms.

Furthermore, our approach did not allow to discriminate between different mechanisms causing variation in tRNA gene content. To obtain a more detailed image of compensatory evolution in *P. aeruginosa*, it will be necessary to analyze the contribution of genomic tRNA genes duplications, tRNA gene acquisition through HGT and mutations in the anticodons of tRNA genes (Yona et al. 2013) to tRNA gene content variation. For example, it would be interesting to know if there are cases of protein coding genes that have been transferred along with tRNA genes that could enhance their expression, similar to what has been observed in viruses (Limor-Waisberg et al. 2011).

Our study shows that codon usage plays an important role in shaping the repertoire of horizontally transferred genes in the *P. aeruginosa* accessory genome and that the incorporation of genes with an atypical codon usage might elicit changes elsewhere in the genome to enhance their integration. The results presented here are however correlative, and there is a need for experiments that quantify the selection coefficient on different synonymous versions of horizontally transferred genes. Experimental manipulation of tRNA gene content will also allow to estimate the fitness effects of horizontally transferred genes in different genomic contexts. Increasing our knowledge on the importance of, and potential evolutionary responses to codon usage in the context of HGT will allow for a better assessment of the potential for the emergence of antibiotic resistance and increased virulence in pathogenic bacteria through HGT.

## Materials & Methods

### Genome sequences

An assembly search was done on ncbi.nlm.nih.gov/assembly (08/09/2017) for “Pseudomonas aeruginosa”. Status = “latest RefSeq” and Assembly level = “complete genome” were selected. This resulted in 96 available full genome assemblies. Two strains were subsequently removed from the dataset. Strain PA7 (GenBank accession GCF_000017205) was removed because it was found to be a taxonomic outlier (see also Roy et al. 2010). Possibly, mutation rates in this strain differ strongly from other strains, making branch length comparison and divergence time with other strains unreliable. Strain H47921 (GenBank accession GCF_001516345) was removed because it was found to be highly recombinant, making a correct phylogenetic placement of this strain difficult. Table S2 provides a list of the genomic sequences included in this study.

Genomes were re-annotated using PROKKA 1.12 (Seemann 2014) to obtain a consistent annotation across all genomes. All subsequent analyses were performed on genes predicted by PROKKA. tRNA genes were predicted using tRNAscan-SE 2.0.5 (Chan et al. 2019).

### Pan-genome analysis

An all-against-all nucleotide BLAST was performed on the full set of genes using blastn 2.2.31+. To cluster genes into groups of homologs, BLAST results were used as input for SiLiX 1.2.9 (Miele et al. 2011) with the following parameter values: overlap (min. % overlap to accept blast hits for building families) = 0.7 and identity (min. % identity to accept blast hits for building families) = 0.6. In parallel, the genome dataset was analyzed with the Roary pipeline (Page et al. 2015) to obtain an additional pan-genome analysis and to use its orthology assignments for splitting up large gene clusters produced by SiLiX.

Our goal here was not to define the *minimal* core genome of *P. aeruginosa*, but to obtain genes for constructing a strain phylogeny and to determine patterns of codon usage for genes expected to be in equilibrium with forces shaping codon usage. Therefore, we handled a looser definition of core genes to maximize the number of genes included in this dataset. This is expected to result in a larger number of core genes compared to other estimates (Freschi et al. 2019). Single copy homologous gene clusters predicted by SiLiX present in at least 95% of all strains (i.e. between 89 and 94 strains) were directly assigned as core genes. Multi-copy clusters predicted by SiLiX present in at least 95% of all strains were split up into single copy clusters using the orthology assignments produced by Roary, and were additionally assigned as core genes.

The remaining homologous gene clusters were assigned as accessory genes. Different reasons could have caused these genes to be classified as accessory genes: HGT, multiple deletions of ancestral core genes, and erroneous classification of some genes in a separate cluster due to, for example, gene duplication followed by sequence evolution leading to pseudogenization or sequencing errors. As only accessory genes that originated through HGT were relevant for our analysis, we filtered out accessory genes that were potentially misclassified or originated through multiple gene deletions. To filter misclassified genes, all accessory gene clusters were blasted against the full set of genes. Only those gene clusters that did not return any significant hits against members of other clusters were retained in the further analysis. Details on filtering accessory genes that potentially originated through multiple deletions of ancestral core genes are described under the section ‘Inference of horizontal gene transfers’.

### Strain phylogeny

All core genes were aligned using MAFFT 7.271 with the ‘auto’ option (Katoh and Standley 2013). A core alignment supermatrix was produced by concatenating all aligned core genes. A maximum likelihood phylogenetic tree was constructed with IQ-Tree 1.6.5 (Nguyen et al. 2014) using a GTR+F+R9 model of substitution chosen according to BIC obtained with ModelFinder (Kalyaanamoorthy et al. 2017). Branch support was obtained with 1000 bootstrap replicates. The strain phylogeny was rooted at midpoint. The position of the root was further supported by several potential EPA placements of *P. pseudoalcaligenes* as outgroup close to the midpoint (Berger et al. 2011).

### Inference of horizontal gene transfers

The overall small number of genes and low amount of sequence variation within homologous clusters of accessory genes did not allow us to construct reliable gene trees, preventing analysis of HGT based on phylogeny comparison (reviewed in Boussau and Scornavacca 2020). Therefore, we used ancestral state reconstruction to infer HGT events. For each accessory gene, we constructed a phyletic pattern with gene copy number as character states. The probability of each character state was inferred for all nodes on the strain phylogeny using ancestral state reconstruction with stochastic character mapping. This was done using the Phytools R package running 1000 simulations for each character under the ER model. We used the probabilities of presence at the nodes of the strain phylogeny to infer HGT events. Horizontal transfer of an accessory gene was predicted to have occurred along a branch on the strain phylogeny if the probability of presence of this gene was lower than 0.5 on its parent node (most probable gene copy number = 0) and higher than 0.5 on its child node (most probable gene copy number > 0). To verify results obtained by Phytools, we additionally inferred HGT events with ancestral reconstruction by posterior probabilities in a phylogenetic birth-and-death model using the COUNT program (Csüös 2010). This analysis inferred the majority of transfers on exactly the same branch (76.7%). Despite some differences between Phytools and COUNT, general patterns in subsequent analyses were highly similar using inferred HGT’s from either program, indicating the robustness of our results. We therefore only included results obtained from HGT’s inferred by Phytools in this study.

Multiple independent gene losses after a single HGT event could produce a scattered phyletic pattern for which multiple false positive HGT events are inferred. To correct for this, we used information on the genomic position of genes for which multiple transfers were inferred. Genes located at the same genomic position (inserted between the same core genes) were assumed to be acquired at the same moment, and their HGT was traced back to the last common ancestor of all separate HGT events inferred for this gene. We removed genes from the analysis that were inferred to be present at the root because their classification as accessory genes seemed to be caused by multiple losses of a gene that was present in the ancestor of all our strains.

Each gene that was acquired through HGT was assigned a ‘residence time’. This residence time was obtained from the strain phylogeny by taking the branch length from the leaf node of the strain containing this gene to the midpoint of the branch where the HGT was inferred. Residence time thus indicates the amount of evolutionary change that occurred in the core genome since a HGT event. We expect this metric to be correlated with actual time, but due to the lack of bacterial fossils, we were unable to fit a molecular clock. Furthermore, it also takes into account the variation in evolutionary rates in the core genome along a branch due to a higher mutation rate or strong selection, something that is also expected to affect the accessory genes contained in the genome.

### Codon usage

The codon usage of the core genome was determined by constructing a codon usage table based on all core genes using the cusp tool (EMBOSS). Metrics regarding codon usage of both core genes and HGT genes were calculated using the COUSIN tool (Bourret et al. 2019) with the codon usage table of the core genome used as a reference. Codon usage metrics were only calculated for genes having at least 100 codons to avoid biases caused by gene length. The Codon Usage Similarity Index (COUSIN; Bourret et al. 2019) was used as a metric to determine how similar the codon usage of a gene was compared to the average codon usage in the core genome. This metric incorporates both the direction of codon usage bias, analogous to the Codon Adaptation Index (CAI; Sharp and Li 1986), and the strength of codon usage bias, analogous to the effective number of codons (ENC; Wright 1990), thus providing more information than each separate metric.

Overall change in COUSIN values for HGT genes in function of residence time was evaluated using a non-parametric Kendall’s rank correlation. Changes in minimal, maximal and median COUSIN values were evaluated by obtaining these values for each residence time interval of 0.001 and fitting a linear model. We additionally analyzed changes in CAI values in function of residence time for comparison. There was a degree of uncertainty if genes inferred to be acquired near the root represented actual HGT events or were ancestral genes with an early loss in one clade (although less probable based on ancestral state reconstruction). Therefore, we also evaluated changes in COUSIN values in function of residence time when excluding HGT genes acquired along branches departing from the root (excluding 159 genes; 6.7% of potential HGT’s).

To evaluate the potential importance of gene amelioration for changes in codon usage in function of residence time, we inferred ancestral sequences of HGT genes using PRANK v.170427 by aligning extant sequences using the codon model and the core genome phylogeny as a guide tree. To allow for a sufficient amount of sequence variation when inferring ancestral sequences, this analysis was only performed on a subset of 22 gene clusters that contained 20 or more extant sequences derived from the same HGT event. The sequence inferred at the ancestral node of the guide tree was used to determine the ancestral codon usage bias. Any gaps in the alignments were removed before assessing their codon usage bias. Significance of change in codon usage bias between the extant genes and their ancestral state was assessed using a one-sample t-test. Genes for whom the mean change in codon usage bias significantly differed from zero were considered to show directional evolution. We additionally assessed the significance of changes in codon usage bias caused only by synonymous variation by converting all non-synonymous mutations back to their ancestral state.

### Evolution of tRNA gene copy numbers

The tRNA gene content was inferred for the parent and child node of each branch on the strain phylogeny using ancestral state reconstruction based on the extant tRNA gene copy numbers found in each strain (same model as for accessory genes). We further estimated the genomic content of protein coding sequences for the parent and child nodes of each branch based on the predicted core genome content and ancestral state reconstruction of HGT genes.

To analyze compensatory evolution on a codon level, codon usage tables were obtained for the parent and child node of each branch based on the inferred genomic content. Changes in codon frequencies due to HGT (expressed as changes in the expected number of a codon per 1000 bases) were calculated by subtracting parent node codon frequencies from child node codon frequencies. A permutation test was used to evaluate if increments in tRNA gene copy number were associated with strong increases in the frequency of codons these tRNAs can translate (McDonald et al. 2015). This was done by ordering changes in codon frequencies of all branches in descending order and assigning a rank number based on their position. The sum of rank numbers of changes in codon frequencies associated with an increase in the tRNA gene having a Watson-Crick base pairing was taken (= observed value). Rank numbers were subsequently randomized using 10^4^ permutations and a p-value was calculated based on the fraction of randomizations for which a rank value sum equal or lower than the observed value was obtained. The same analysis was performed separately for codons associated with increases in tRNA genes having wobble- and less-favored wobble base pairing (Murphy and Ramakrishnan 2004).

To analyze compensatory evolution on a gene level, we first calculated the absolute adaptiveness (dos Reis et al. 2004) of each codon for the parent and child nodes of branches along which a change in tRNA gene content was observed. The absolute adaptiveness of a codon i, denoted as W_*r*_ was defined as:

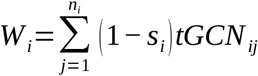

with n_*i*_ being the number of tRNA isoacceptors that recognize the *i*th codon, tGCN_*ij*_ the gene copy number of the *j*th tRNA that recognizes the *i*th codon, and s_*ij*_ a selective constraint on the efficiency of the codon-anticodon pairing. As no experimental data is available for estimating the values of s_*ij*_ in *P. aeruginosa*, we used s_*ij*_ = 0 for Watson-Crick pairing, s_*ij*_ = 0.5 for wobble pairing and s_*ij*_ = 0.9 for less favored wobble pairing. The expected translation efficiency of a gene in a given tRNA context is then calculated as the geometric mean of the absolute adaptiveness of its codons. This metric is highly similar to the tAI (dos Reis et al. 2004), which was designed to compare different genes to the same tRNA pool and uses the relative adaptiveness of codons instead of the absolute adaptiveness. However, tAI was not directly applicable to our question of comparing the translation efficiency of the same genes in the context of a variable tRNA pool because duplications in already favored tRNA isoacceptors caused either no change or a lowering of the tAI value of all genes, which is biologically not meaningful. Using the absolute adaptiveness instead of relative adaptiveness of codons solved this issue. Because of its high similarity to tAI, we will refer to this metric as tAI_abs_. Change in the tAI_abs_ of each gene was calculated by subtracting the tAI_abs_ obtained with the tRNA gene content at the parent node from the tAI_abs_ obtained with the tRNA gene content at the child node (ΔtAI_abs_). For each branch where the tRNA gene content variation caused a positive mean ΔtAI_abs_, a non-parametric Mann-Whitney U-test was used to evaluate if the ΔtAI_abs_ differed significantly between HGT genes acquired along that branch and genes already present at the parent node.

## Acknowledgments

This work was supported by the European Research Council (Consolidator Grant HGTCODONUSE number 682819). The authors would like to thank Léa Pradier, Caroline Rose, Enrique Ortega-Abboud, Michael Finnegan, Marie-Pierre Dubois, Laurent Duret and members of the Génétique et Ecologie Evolutive team for stimulating discussions and constructive comments on the manuscript.

