## Supplementary Figures for "Evolutionary responses to codon usage of horizontally transferred genes in *Pseudomonas aeruginosa*"

### Figure S1

**Schematic overview of the bioinformatic pipeline used in this study**. A: pan-genome analysis; B. strain phylogeny; C. Filter accessory genes. D. Ancestral state reconstruction and inference of HGT's

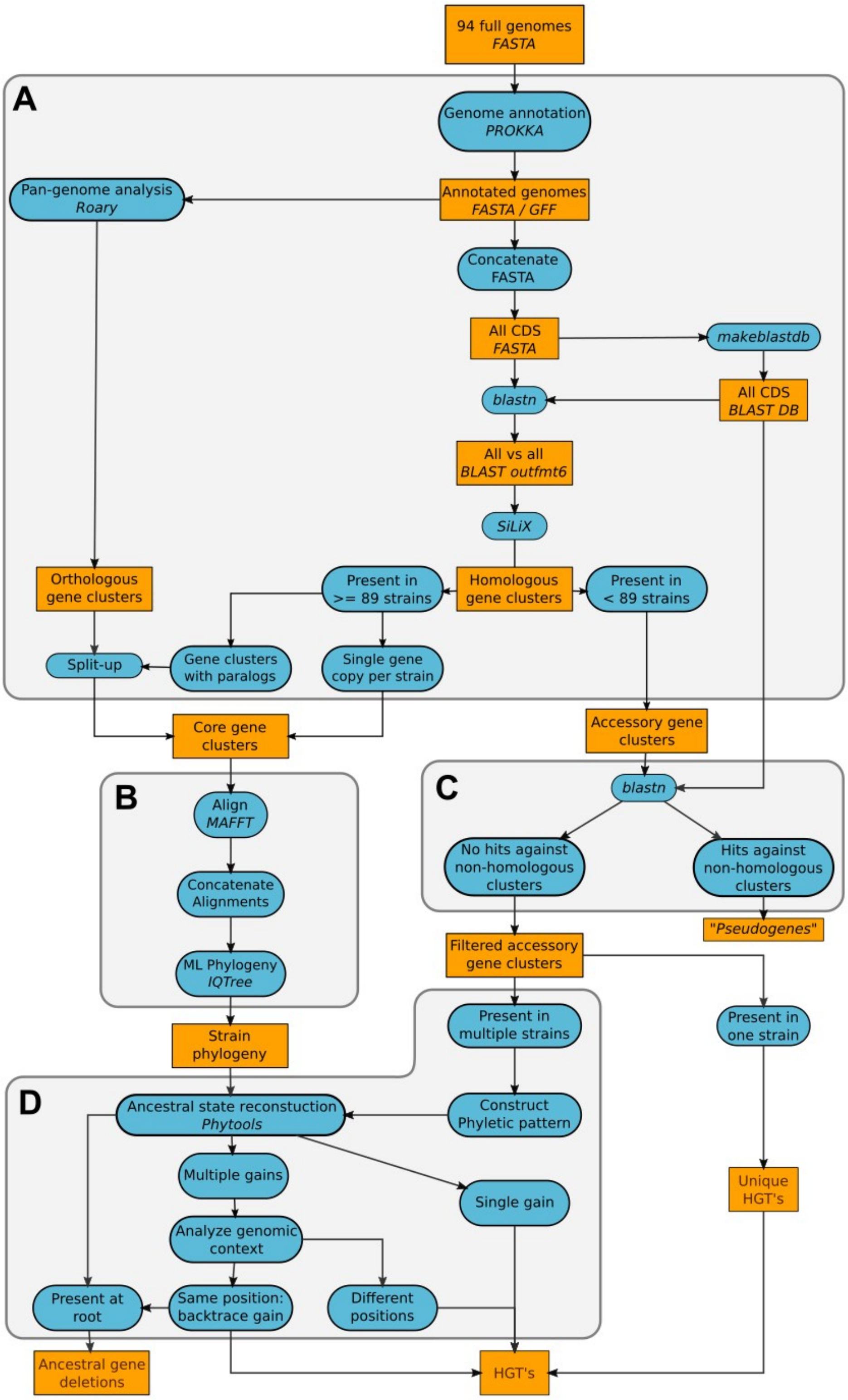

### Figure S2

**Phylogenetic tree containing the *P. aeruginosa* strains used in this study**. The tree was produced using the concatenated alignment of all the core genes and inferred using a GTR+F+R9 model of substitution. The tree was rooted at midpoint. Branch support was obtained with 1000 bootstrap replicates. Bootstrap values are indicated on the nodes.

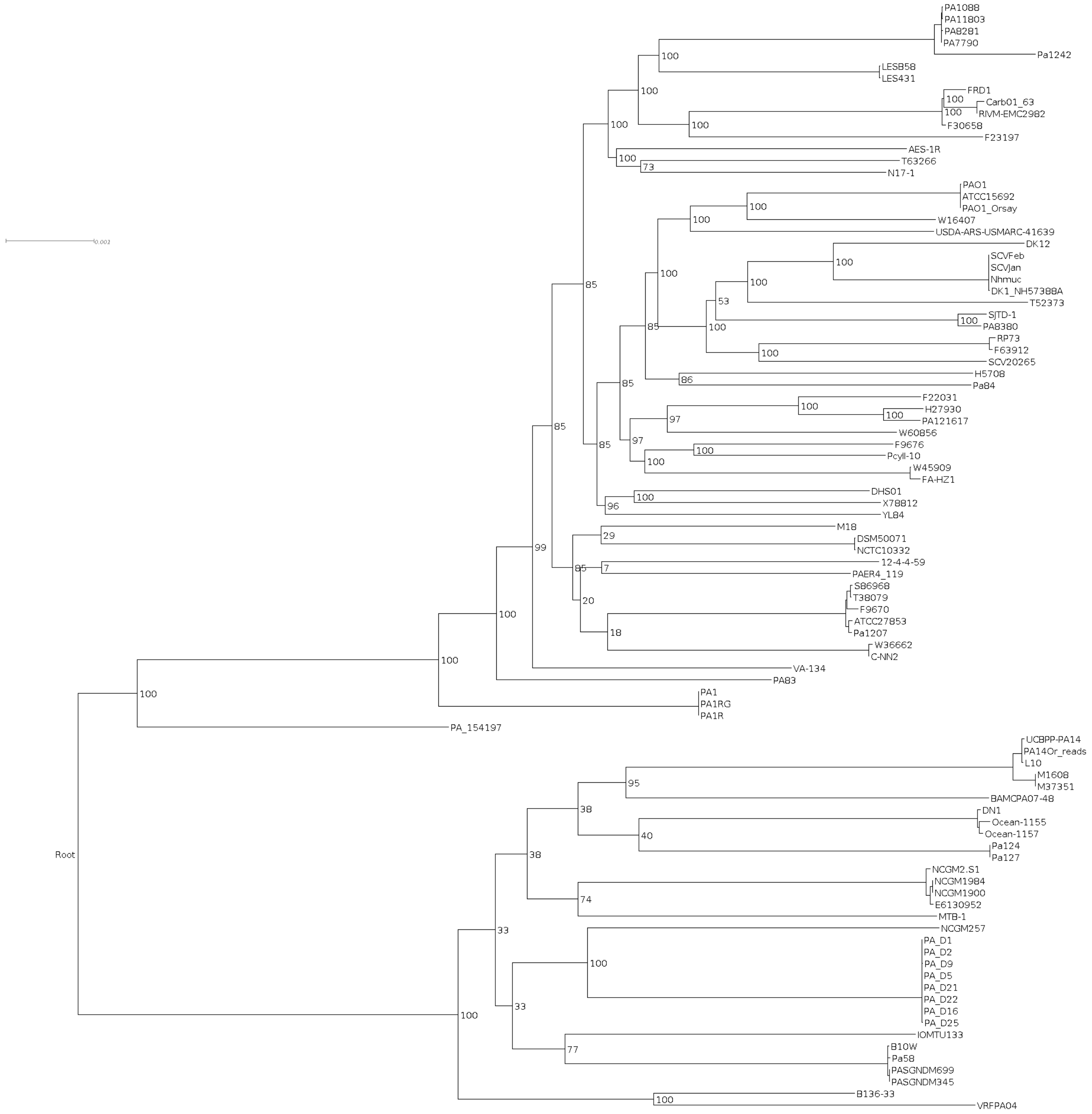

### Figure S3

**Distribution of the residence time of genes acquired by HGT.** The largest fraction of genes were recently acquired, while at longer residence times the amount of genes was strongly reduced. The inset shows the distribution of distances between the tree tips and all internal nodes of the strain phylogeny, indicating that the gaps in the distribution of residence time are potentially caused by the structure of the strain phylogeny.

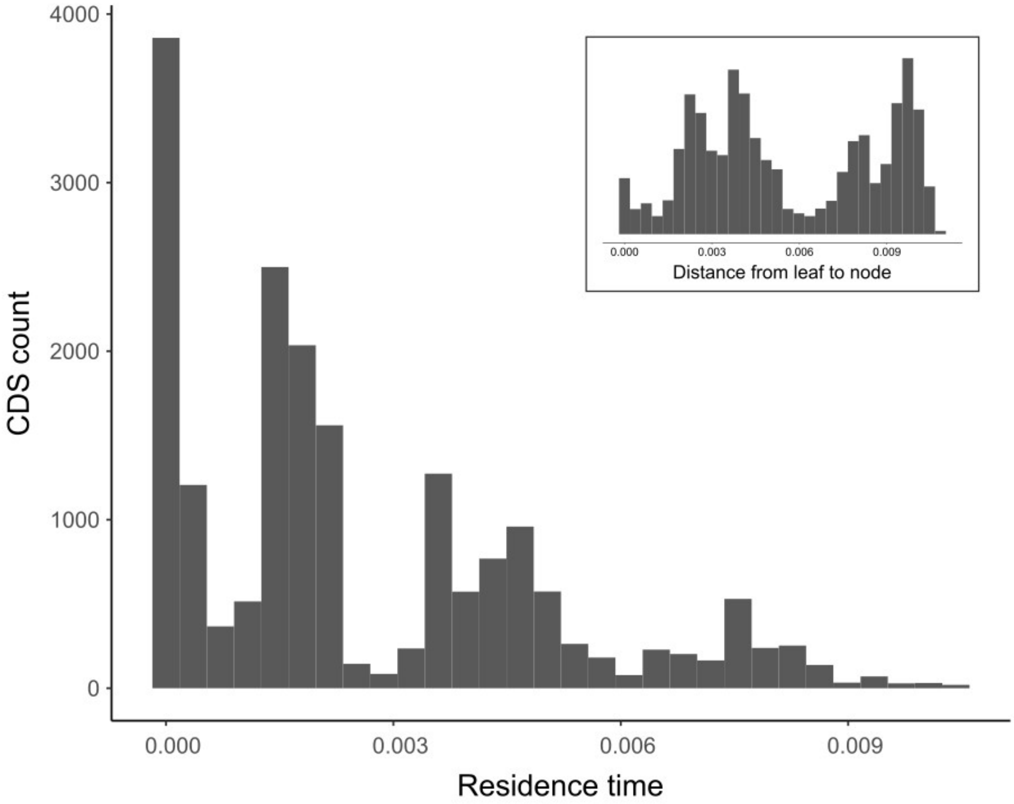

### Figure S4

**Codon usage bias of HGT genes in relation to their residence time.** Scatterplot of the CAI value of HGT genes in function of their residence time. The blue and red regression lines were determined by taking respectively the minimal and maximal CAI value for each residence time interval of 0.001.

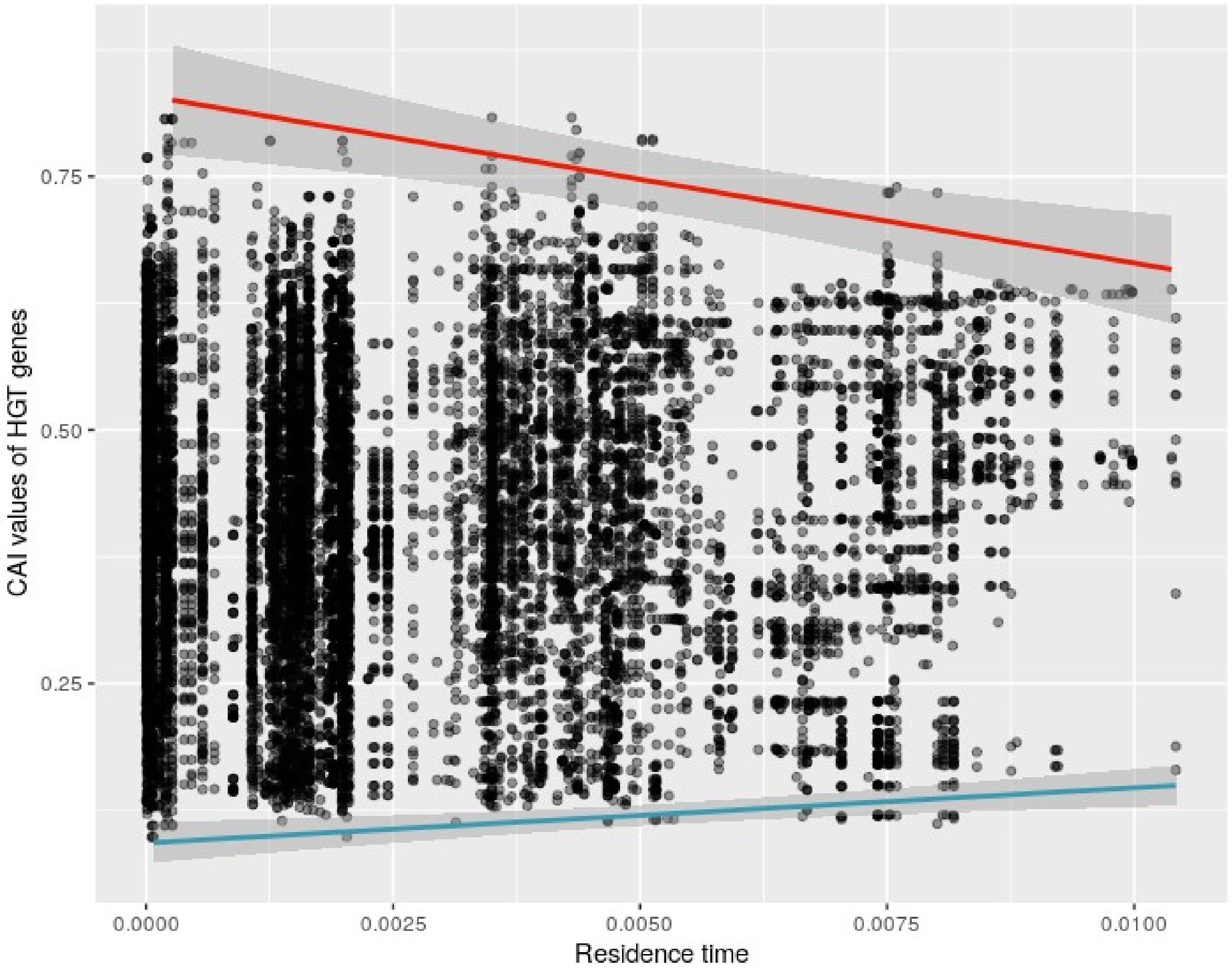

### Figure S5

**Variation in tRNA gene copy number in *P. aeruginosa*.** Each tRNA gene is specified by its anticodon, isoacceptor tRNA's are grouped by amino acid. The y-axis indicates the copy number that was observed for a specific tRNA gene, the circle size indicates the relative fraction of strains having a specific tRNA gene copy number.

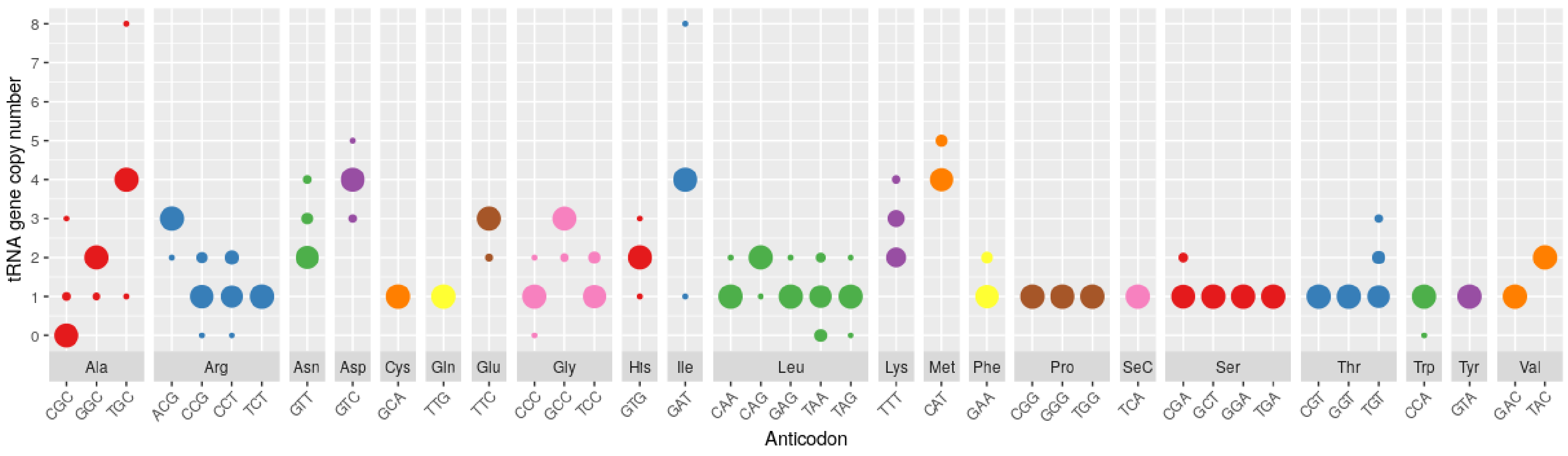
